# Satiation is associated with OGT-dependent regulation of excitatory synapses

**DOI:** 10.1101/2025.09.02.673689

**Authors:** Mario Pérez Del Pozo, Manish Bhattacharjee, Anushree Tripathi, Thyra Boafo, Sabrina Galizia, Paolo Medini, Michael Druzin, Olof Lagerlöf

## Abstract

Satiation is essential for energy homeostasis and is dysregulated in metabolic disorders like obesity and eating disorders such as anorexia nervosa. While satiation engages a large neural network across brain regions, how the communication within this network depends on metabolic fluctuations is unclear. This study shows that nutrient access can affect neuron-to-neuron communication in this network by regulating excitatory synaptic plasticity through O-GlcNAc transferase (OGT) in αCaMKII satiation neurons in the paraventricular nucleus (PVN). Using cell-specific knockout mice and electrophysiological recordings, we demonstrate that OGT deletion in PVN^αCaMKII^ neurons increases input resistance and neuronal excitability while preserving basic membrane electrical properties. Strikingly, feeding triggered a robust 3.8-fold increase in the excitatory synaptic input in wild-type neurons, whereas OGT-knockout neurons failed to exhibit this feeding-induced synaptic activation, instead displayed a paradoxical trend towards increases in synaptic activity during hungry conditions. Furthermore, OGT deletion destabilized glucose-dependent synaptic responses, with knockout neurons displaying maladaptive depression of excitatory transmission in conditions where stability is normally preserved. These findings establish OGT as a nutrient-sensitive modulator of synaptic plasticity that ensures appropriate satiation signalling by coupling metabolic state to synaptic plasticity.

## Introduction

Satiation is the process that occurs during eating and leads to meal termination, providing the physiological signal to stop consuming the current meal. It is a fundamental and complex physiological process critical for energy homeostasis. It is regulated by a network of neural and hormonal signals [1, 2]. Many of these signals affect food intake through the hypothalamus in the brain. The paraventricular nucleus (PVN) of the hypothalamus plays a fundamental role for feeding behavior by integrating peripheral signals of energy status with emotional and cognitive aspects of feeding behavior [3-5]. Some of its neurons communicate with other parts of the brain through direct neuron-to-neuron contacts, or synapses, while other neurons are part of the neuroendocrine system and affect body metabolism through sending processes to the pituitary and median eminence. Disruption of these mechanisms can lead to dysregulated eating patterns and contribute to metabolic disorders such as obesity as well as eating disorders including anorexia nervosa [6], highlighting the importance of understanding how their function is coupled to the metabolic state of the body.

Central to this metabolic coupling is the brain’s ability to monitor glucose availability through specialized glucose-sensing mechanisms within hypothalamic circuits[7, 8].

The most well-characterized system is the classical GLUT2/glucokinase/K_ATP_ channel pathway that creates glucose-excited and glucose-inhibited neurons analogous to pancreatic β-cells[9-11]. These threshold-based sensors respond to glucose concentrations within their physiological range and have proven essential for basic glucose homeostasis, particularly in counterregulatory responses to hypoglycemia. They are either turned on or turned off, respectively, by glucose and in response regulate intrinsic excitability of the neuron.

In addition to these established pathways, a significant fraction of nutrients taken up by cells is metabolized through the hexosamine biosynthesis pathway (HBP) to uridine-diphosphate N-acetylglucosamine (UDP-GlcNAc). UDP-GlcNAc can then be cleaved by the enzyme O-GlcNAc transferase (OGT). OGT catalyzes the transfer of β-N-acetylglucosamine (GlcNAc) to serine and threonine residues of intracellular proteins (O-GlcNAc)[12]. O-GlcNAc can then be removed by O-GlcNAcase (OGA). Its dependence on flux through the HBP and regulation by other metabolic pathways such as insulin suggest that OGT regulates cellular function depending on the body’s metabolic status[13]. Indeed, the HBP and O-GlcNAc cycling are linked genetically to body weight and metabolic disorders in humans[13-16] We have previously shown that OGT can regulate food intake. In the PVN, O-GlcNAc levels are sensitive to metabolic fluctuations particularly in αCaMKII-expressing cells (PVN^αCaMKII^)[17] While these neurons have not been fully characterized, our previous observations show that these neurons become activated upon food intake. Once activated, as revealed by optogenetics, they turn off further intake[5]. This feeding-induced activation is completely dependent on OGT. Deleting OGT in these neurons during a time course where there is no apparent effect on cell health increases food intake and body weight rapidly due to larger meal size. It may be the case that this effect on energy homeostasis is compensated by other neurons after several months[17-20]. Furthermore, αCaMKII-expressing neurons in certain brain regions have been shown to suppress further food intake upon activation, as demonstrated in the anterior insular cortex, pointing to a broader role for these neurons in feeding regulation.[21] These and other data indicate that the PVN^αCaMKII^ neurons mediate satiation, at least in part through OGT. We and others have reported that OGT can regulate neuron function by affecting excitatory neurotransmission: OGT regulates both excitatory synapse number and synapse strength in part by controlling the synaptic abundance of AMPA receptors[17, 22] It has been shown also that glutamatergic neurotransmission through the AMPA receptor in the PVN regulates food intake[23, 24]. However, unlike classical glucose sensors that exhibit binary responses to acute nutrient changes, it is unclear whether and how OGT may affect excitatory neurotransmission in PVN^αCaMKII^ depending on nutrient access to affect satiation.

The current study investigates the electrophysiological mechanisms through which OGT regulates PVN^αCaMKII^ neuronal function under varying metabolic conditions. We demonstrate that selective deletion of OGT from PVN^αCaMKII^ increases neuronal input resistance and increases excitability while preserving basic membrane electrical properties. Most strikingly, we identify a critical role for OGT in coupling nutritional status to synaptic transmission, as OGT-knockout (OGT-KO) neurons exhibit a paradoxical decline in excitatory synaptic input during feeding—the opposite pattern observed in wild-type (WT) neurons, where feeding strongly enhances excitatory synaptic input. Furthermore, we demonstrate that OGT maintains robust synaptic responses during acute glucose fluctuations, with OGT-KO neurons displaying marked synaptic depression when exposed to elevated glucose. Taken together, these findings indicate that OGT couples nutrient availability to satiation by regulating excitatory synaptic plasticity in PVN^αCaMKII^.

## Results

### PVN^αCaMKII^ are parvocellular neurons

While the PVN^αCaMKII^ neurons are critical for satiation, their basic morphological and electrophysiological properties have not been characterized. To selectively label and delete OGT from PVN^αCaMKII^ neurons, we utilized stereotactic injections of a cocktail of two AAV1 vectors encoding CaMKII-Cre and floxed enhanced green fluorescent protein (EGFP), respectively, into the PVN of both OGT-floxed and wildtype (WT) C57BL/6 mice (Figure 1A). Immunohistochemical analysis via confocal microscopy confirmed robust transduction as evidenced by intense EGFP expression localized within the PVN. We used low-titer viral stocks to achieve sparse OGT deletion in a limited population of PVN^αCaMKII^ neurons. This preserves overall network function and ensures that subsequent electrophysiological analyses reflect cell-autonomous effects in the targeted neurons rather than secondary effects from hyperphagia-dependent obesity. OGT immunoreactivity was largely absent in EGFP-labeled KO-neurons, yet residual OGT signal remained in other cells in the PVN, reflecting our deliberately sparse knockout strategy designed to isolate cell-autonomous effects without inducing secondary systemic changes (Figure 1B). We confirmed whether sparse OGT deletion influenced systemic parameters: there was no significant difference in average body weight between WT and OGT-KO mice (WT: 25.3 ± 1.11 g; OGT-KO: 25.37 ± 0.54 g; P = 0.956) (Figure 1C), nor in daily food intake (WT: 4.88 ± 0.29 g; OGT-KO: 4.94 ± 0.53 g; P = 0.925) (Figure 1D). Our previous observations show that PVN^αCaMKII^ overlap with markers for both thyrotropin-releasing hormone (TRH) and oxytocin neurons[17]. TRH is specifically produced by parvocellular neurons[25], while oxytocin can be produced by both parvocellular and magnocellular neurons in the PVN[26, 27]. However, quantitative morphometric analysis of 153 EGFP-positive neurons revealed a mean cell diameter of 10.79 µm (±0.206 µm SEM; standard deviation, 1.842 µm), consistent with the expected size distribution of only parvocellular neurons (Figure 1E). Parvocellular neurons that express a low-threshold spike (LTS) typically send axons to other parts of the brain rather than directly regulating peripheral metabolism through the neurosecretory system in the pituitary[28]. A substantial fraction of PVN^αCaMKII^ neurons (6/16; 37.5%) exhibit low-threshold spikes(LTS) (Figure 1F, G), marking this as the first report of LTS in this genetically defined population in the PVN. These findings indicate that PVN^αCaMKII^ mediate satiation at least in part through participating in a larger intra-brain neural network.

**Figure 1.**
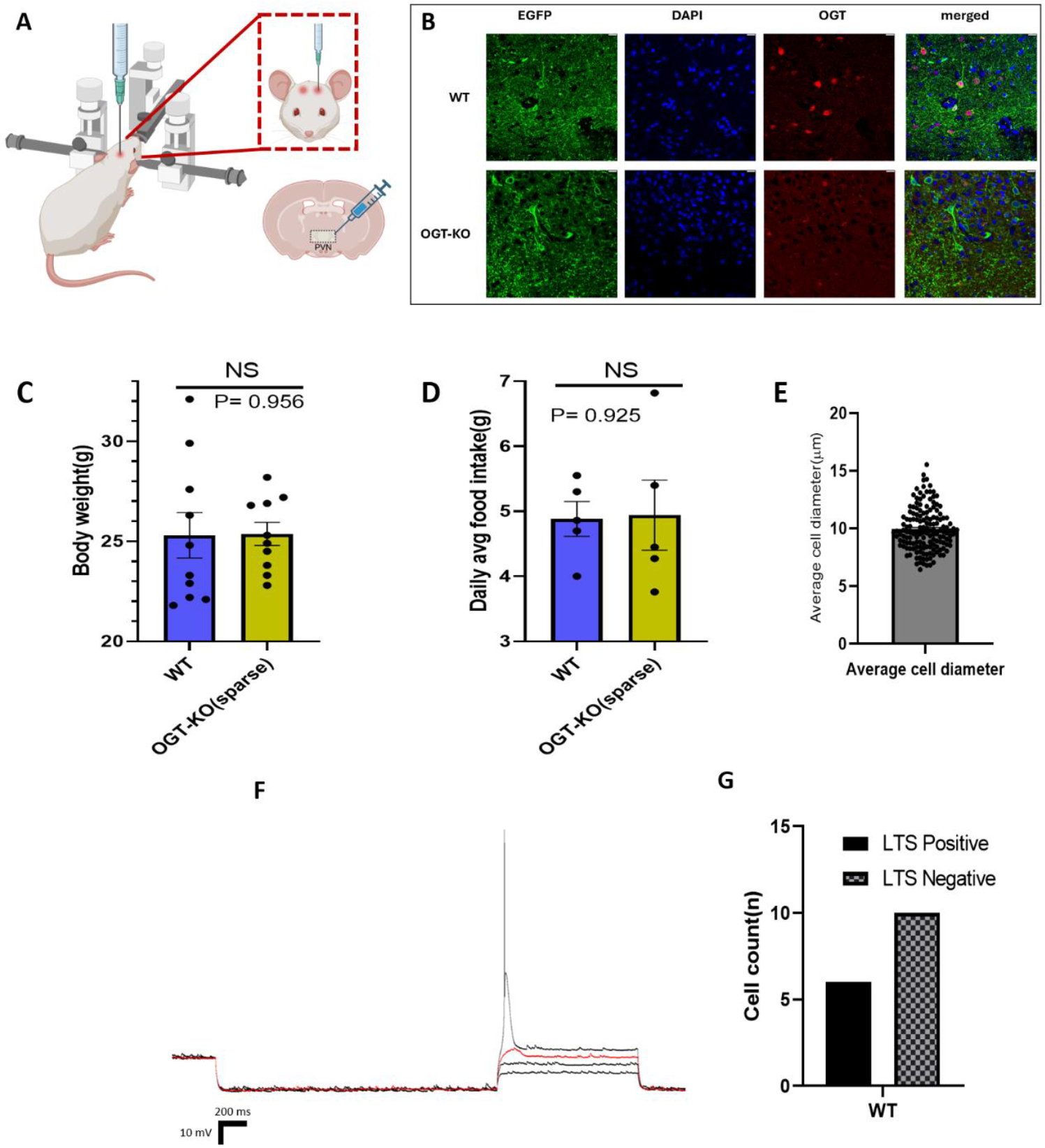
OGT-KO in the PVN^αCaMKII^ Predominantly Targets Parvocellular Neurons: **(A)** Schematic of stereotactic injection procedure in PVN of OGT-floxed and WT C57BL/6 mice. **(B)** Representative confocal micrographs showing EGFP expression (green) and OGT immunolabeling (red) in WT and OGT-KO brain slices. DAPI counterstains nuclei (blue). Scale bar: 10 µm. **(C)** Average body weight over 10-day post-surgical period comparing WT vs OGT-KO mice (NS, P = 0.956). **(D)** Daily average food intake over same period comparing genotypes (NS, P = 0.925). **(E)** Cell diameter analysis of EGFP-expressing neurons showing mean diameter of 10.79 µm (±0.206 µm SEM; SD = 1.854 µm). **(F)** Representative current-clamp trace showing Low-threshold spike (LTS) detection protocol in WT neurons. The red sample trace is representative of the LTS; the red arrow head points at the tiny spike observed characteristic feature of LTS **(G)** LTS classification showing 6/16 (37.5%) LTS-positive and 10/16 (62.5%) LTS-negative WT neurons. Data from OGT-floxed (n = 10) and WT (n = 10) mice, except panel E where OGT-floxed (n = 5) and WT (n = 5) mice. Cell diameter measured from 153 EGFP-expressing cells. Statistical comparisons using Mann-Whitney test. Data presented as mean ± SEM. Blue bars: WT; yellow bars: OGT-KO.

### OGT-KO in the PVN^αCaMKII^ does not alter basic electrophysiological properties but increases neuronal excitability

Basic electrophysiological properties were measured to determine whether OGT deletion affects the intrinsic biophysical characteristics of PVN^αCaMKII^ cells neurons. Representative traces from WT and OGT-KO neurons during AP threshold and baseline membrane potential measurements are shown in Figures 2A and 2B, respectively. Resting membrane potential (WT: −74.02 ± 14.68 mV; OGT-KO: −65.52 ± 18.09 mV; P = 0.226), action potential(AP) threshold (WT: −41.32 ± 10.87 mV; OGT-KO: −45.31 ± 16.42 mV; P = 0.226), and membrane capacitance (WT: 52.77 ± 34.29 pF; OGT-KO: 40.38 ± 17.77 pF; P = 0.368) were not significantly different between WT and OGT-KO neurons (Figures 2 C-E). These observations show that deleting OGT does not affect basic neuron properties in PVN^αCaMKII^.

**Figure 2.**
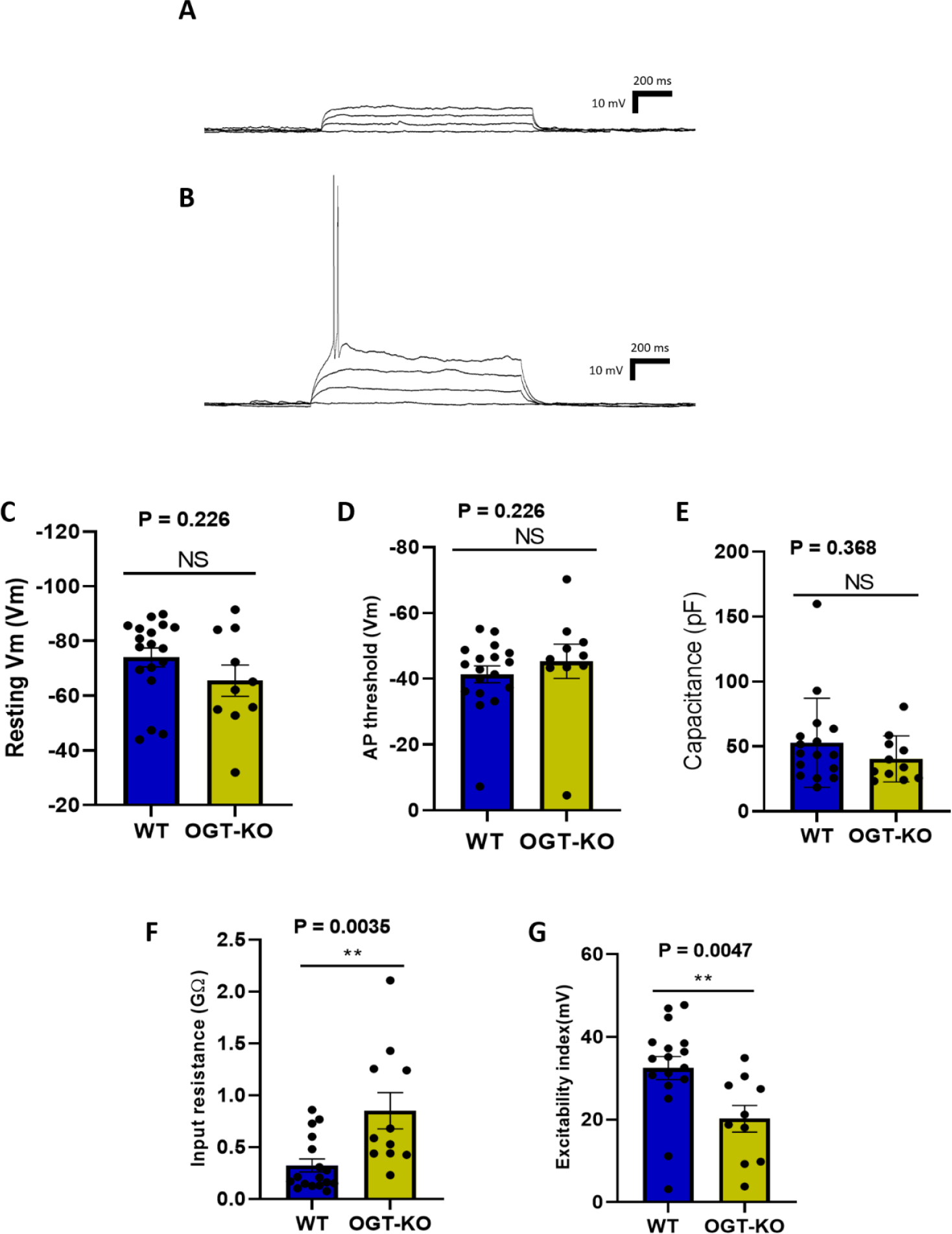
OGT-KO in the PVN^αCaMKII^ alters input resistance and neuronal excitability without altering basic electrophysiological properties: Representative current-clamp traces for action potential threshold and baseline membrane potential measurements in **(A)**WT and **(B)**OGT-KO neurons. **(C)** Resting membrane potential comparing WT vs OGT-KO neurons (NS, P = 0.226). **(D)** Action potential threshold comparing genotypes (NS, P = 0.226). **(E)** Membrane capacitance comparing genotypes (NS, P = 0.368). **(F)** Input resistance comparing WT vs OGT-KO neurons (P = 0.0035). **(G)** Excitability index comparing genotypes (P = 0.0047). Data from WT (n = 17-18) and OGT-KO (n = 10-11) PVN parvocellular neurons. Statistical comparisons using Mann-Whitney non-parametric test. Data presented as mean ± SEM. Blue bars: WT neurons; yellow bars: OGT-KO neurons.

In contrast, properties related to neuronal responsiveness were significantly altered in OGT-KO neurons. Input resistance was markedly increased in OGT-KO neurons (WT: 0.32 ± 0.26 GΩ; OGT-KO: 0.85 ± 0.58 GΩ; P = 0.0035) (Figure 2F). The increase in input resistance in OGT-KO neurons implies that even small current inputs will result in larger changes in membrane voltage, effectively amplifying synaptic input. To further characterize the functional consequences of altered input resistance on neuronal excitability, we next examined the threshold for action potential generation by measuring the excitability index. The excitability index is defined as the minimum depolarization required to elicit an AP. Consistent with the increased input resistance, OGT-KO neurons showed a significantly lower excitability index compared to WT neurons (WT: 32.47 ± 11.49 mV; OGT-KO: 20.21 ± 10.23 mV; P = 0.0047) (Figure 2G), meaning they required less current injection to fire action potentials and were therefore more excitable [29]. These complementary findings demonstrate that OGT deletion enhances the ability of neurons to generate action potentials in response to smaller stimuli, indicating that OGT regulates neuronal excitability in PVN^αCaMKII^ [30].

### OGT regulates feeding-dependent changes in excitatory synaptic input

Our previous work showed that O-GlcNAc levels in PVN^αCaMKII^ cells decrease during fasting and that OGT regulates excitatory synaptic function[17, 24]. To reveal how OGT influences the response of these neurons to different nutritional states, we recorded spontaneous excitatory postsynaptic currents (sEPSCs) from WT and OGT-KO neurons obtained from mice under hungry and fed conditions (Figure 3).

**Figure 3.**
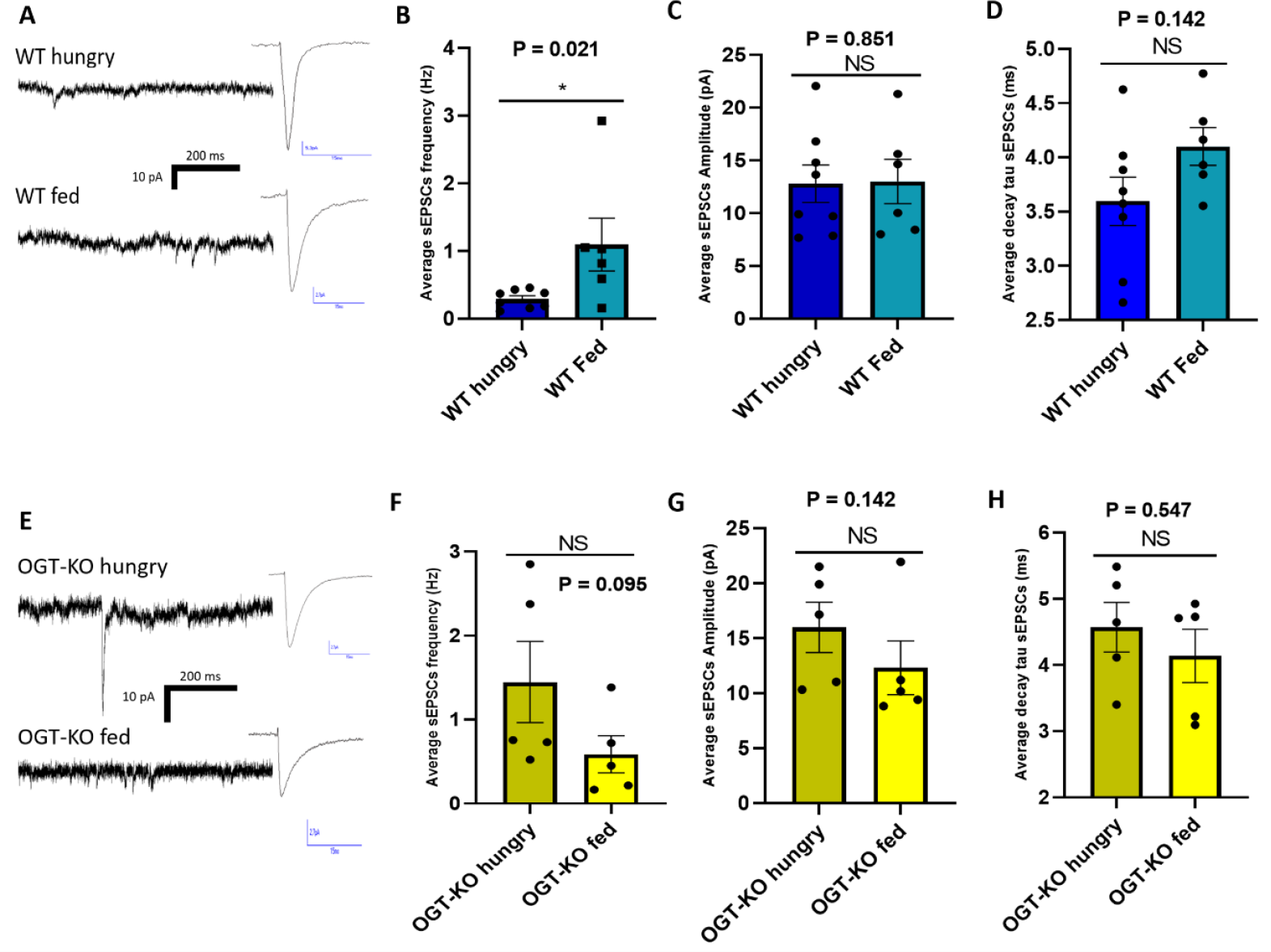
Nutritional State Differentially Modulates Synaptic Transmission in WT and OGT-KO Neurons: **(A)** Representative sEPSC traces in WT neurons under hungry (top) and fed (bottom) conditions. **(B)** sEPSC frequency in WT neurons comparing hungry vs fed states (P = 0.021). **(C)** sEPSC amplitude in WT neurons comparing hungry vs fed states (NS, P = 0.851). **(D)** sEPSC decay tau in WT neurons comparing hungry vs fed states (NS, P = 0.142b **(E)** Representative sEPSC traces in OGT-KO neurons under hungry (top) and fed (bottom) conditions. **(F)** sEPSC frequency in OGT-KO neurons comparing hungry vs fed states (NS, P = 0.095). **(G)** sEPSC amplitude in OGT-KO neurons comparing hungry vs fed states (NS, P = 0.309). **(H)** sEPSC decay tau in OGT-KO neurons comparing hungry vs fed states (NS, P = 0.547). Data from WT (n = 5) and OGT-KO (n = 5) mice under hungry and fed conditions (n = 5 each condition). Statistical comparisons using Mann-Whitney non-parametric test. Data presented as mean ± SEM. Blue bars: WT neurons (dark blue = hungry, light blue = fed); yellow bars: OGT-KO neurons (light yellow = hungry, bright yellow = fed).

In WT neurons from fed mice, sEPSC frequency was robustly elevated (1.10 ± 0.95 Hz) compared to neurons from hungry mice (0.29 ± 0.13 Hz; P = 0.021) (Figure 3B). This demonstrates that feeding activates excitatory input to PVN^αCaMKII^ neurons, with fed animals showing a nearly 3.8-fold increase in synaptic drive that supports satiation signaling and meal termination.

In contrast, OGT-KO neurons exhibited an opposite trend, with feeding resulting in more than a two-fold lower sEPSC frequency than in the hungry state, although this difference did not reach statistical significance (OGT-KO hungry: 1.45 ± 1.08 Hz; OGT-KO fed: 0.59 ± 0.50 Hz; P = 0.095) (Figure 3F).

The amplitude of sEPSCs was not significantly affected by nutritional state in either WT (WT hungry: 12.81 ± 5.00 pA; WT fed: 13.01 ± 5.16 pA; P = 0.851) or OGT-KO neurons (OGT-KO hungry: 15.98 ± 5.10 pA; OGT-KO fed: 12.31 ± 5.46 pA; P = 0.309) (Figures 3C and 3G). Similarly, the decay tau of sEPSCs remained consistent across nutritional states in both genotypes (WT hungry: 3.59 ± 0.63 ms; WT fed: 4.10 ± 0.43 ms; P = 0.142; OGT-KO hungry: 4.57 ± 0.84 ms; OGT-KO fed: 4.14 ± 0.90 ms; P = 0.547) (Figures 3D and 3H), indicating that OGT specifically regulates synaptic event frequency rather than strength over periods of longer nutrient fluctuations.

These results suggest that OGT plays a critical role in regulating the frequency of excitatory synaptic input to PVN^αCaMKII^ neurons in response to feeding state, while not affecting event amplitude or kinetics. In WT neurons, excitatory drive increases upon feeding, consistent with a satiety-promoting mechanism[17]. In contrast, OGT-KO neurons have higher excitatory input during fasting and lack feeding-induced enhancement, indicating that OGT is necessary for proper state-dependent modulation of synaptic activity.

### OGT stabilizes synaptic responses to acute glucose fluctuations

Since OGT activity depends on UDP-GlcNAc derived from glucose metabolism and is implicated in nutrient sensing and feeding regulation[31]—we hypothesized that OGT may regulate neuronal responses to acute changes in glucose levels during a time course similar to postprandial changes in glucose concentration.

To directly examine how OGT mediates the neuronal response to glucose availability, we exposed acute brain slices to sequential glucose concentrations of 2.5 mM followed by 12.5 mM (Figure 4).

**Figure 4.**
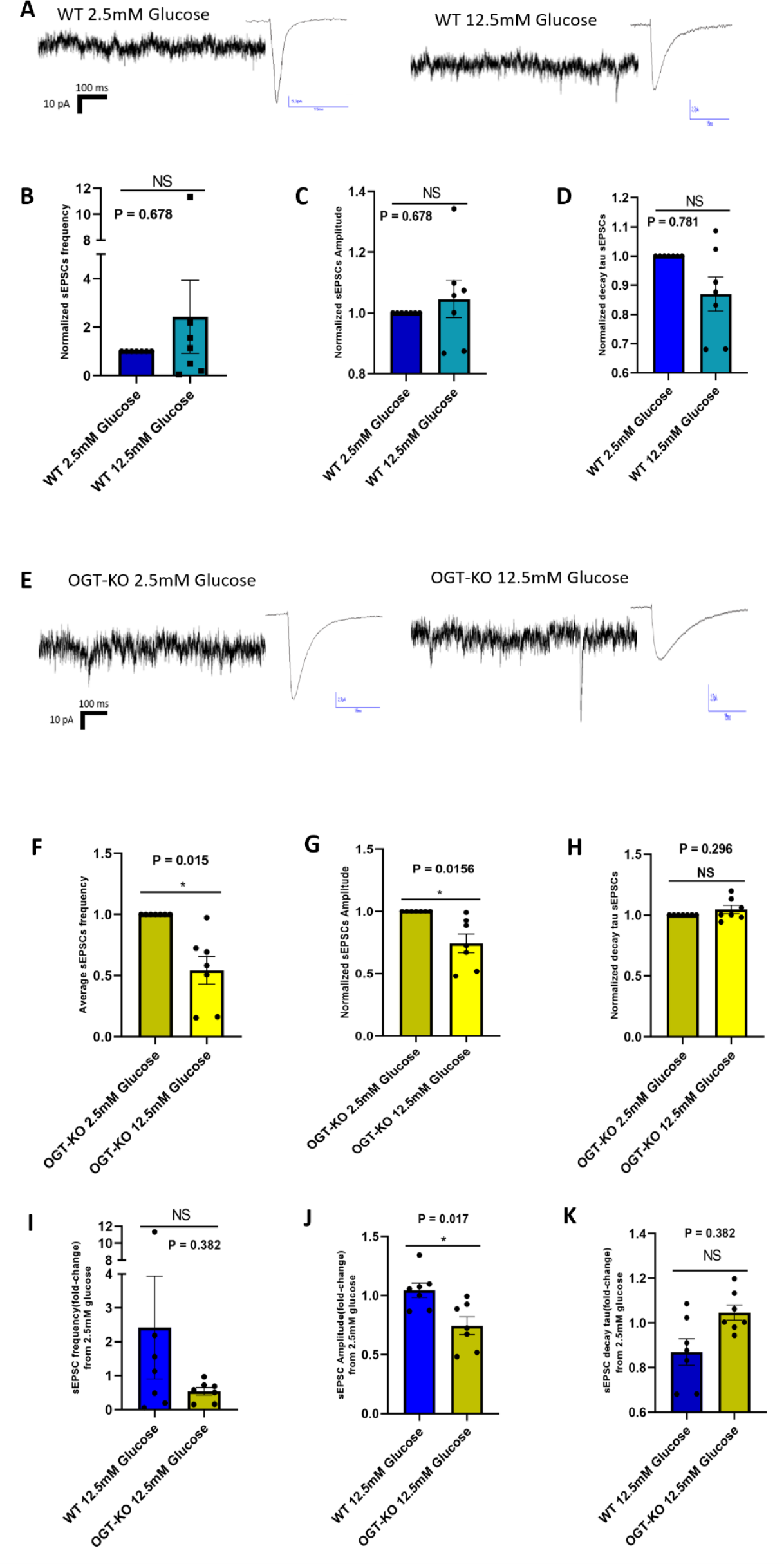
OGT stabilizes synaptic responses to acute glucose fluctuations: **(A)** Representative sEPSC traces in WT neurons under 2.5 mM glucose (left) and 12.5 mM glucose (right). **(B)** Normalized sEPSC frequency in WT neurons comparing 2.5 mM to 12.5 mM glucose (NS, P = 0.687). **(C)** Normalized sEPSC amplitude in WT neurons comparing 2.5 mM to 12.5 mM glucose (NS, P = 0.687). **(D)** Normalized sEPSC decay tau in WT neurons comparing 2.5 mM to 12.5 mM glucose (NS, P = 0.0781). **(E)** Representative sEPSC traces in OGT-KO neurons under 2.5 mM glucose (left) and 12.5 mM glucose (right). **(F)** Normalized sEPSC frequency in OGT-KO neurons comparing 2.5 mM to 12.5 mM glucose (P = 0.0156). **(G)** Normalized sEPSC amplitude in OGT-KO neurons comparing 2.5 mM to 12.5 mM glucose (P = 0.0156). **(H)** Normalized sEPSC decay tau in OGT-KO neurons comparing 2.5 mM to 12.5 mM glucose (NS, P = 0.296). Fold-change analyses (12.5 mM relative to 2.5 mM glucose): **(I)** sEPSC frequency fold-change between WT and OGT-KO neurons (NS, P = 0.382). **(J)** sEPSC amplitude fold-change between WT and OGT-KO neurons (P = 0.017). **(K)** sEPSC decay tau fold-change between WT and OGT-KO neurons (NS, P = 0.382). Data from WT (n=7) and OGT-KO (n=7) mice. Normalized values in B-H set 2.5 mM glucose as baseline (=1). Panels B-H analyzed using Wilcoxon signed-rank test; panels I-K analyzed using Mann-Whitney test. Data presented as mean ± SEM. Blue bars: WT neurons; yellow bars: OGT-KO neurons.

In WT neurons, acute elevation of glucose from 2.5 mM to 12.5 mM did not significantly affect normalized sEPSC frequency (12.5 mM Glucose = 2.42 ± 4.0; P = 0.687) (Figure 4B), amplitude (12.5 mM Glucose = 1.05 ± 0.16; P = 0.687) (Figure 4C), or decay tau (12.5 mM Glucose = 0.87 ± 0.16; P = 0.0781) (Figure 4D). This suggests that WT PVN^αCaMKII^ neurons maintain stable synaptic properties during acute glucose fluctuations[18].

In marked contrast, OGT-KO neurons exhibited a significant decrease in synaptic transmission when exposed to elevated glucose. sEPSC frequency markedly decreased in OGT-KO neurons at 12.5 mM glucose (12.5 mM Glucose = 0.54 ± 0.3; P = 0.0156) (Figure 4F). Similarly, sEPSC amplitude was significantly reduced in OGT-KO neurons under high glucose conditions (12.5 mM Glucose = 0.74 ± 0.20; P = 0.0156) (Figure 4G), while decay tau remained unchanged (12.5 mM Glucose = 1.05 ± 0.09; P = 0.296) (Figure 4H). This differential response to acute glucose fluctuations provides evidence that OGT functions to stabilize excitatory synaptic input during metabolic fluctuations, adapting the PVN^αCaMKII^ response to glucose elevations rather than being necessary to detect changes in glucose fluctuations.

When comparing the fold-change responses (12.5 mM Glucose/2.5 mM Glucose) between WT and OGT-KO neurons, we observed that the frequency response did not reach statistical significance (WT: 2.42 ± 4.00; OGT-KO: 0.54 ± 0.30; P = 0.382) (Figure 4I), likely due to high variability in WT neurons. However, the amplitude response was significantly different (WT: 1.05 ± 0.16; OGT-KO: 0.74 ± 0.20; P = 0.0175) (Figure 4J). The comparison of decay tau fold-changes between the groups showed a trend toward significance (WT: 0.87 ± 0.16; OGT-KO: 1.05 ± 0.09; P = 0.053), although it did not reach the conventional threshold for statistical significance (Figure 4K). These results further substantiate that OGT regulates glucose-dependent synaptic plasticity in PVN^αCaMKII^ neurons.

These findings favor a model in which OGT regulates excitatory synaptic input to PVN^αCaMKII^ neurons in response to metabolic fluctuations. In wild-type neurons, excitatory input appropriately increases upon feeding and support satiation, whereas OGT-KO neurons exhibit paradoxical synaptic responses to metabolic fluctuations[32] (Figure 5). Together, our data show that OGT adapts synaptic plasticity to nutritional status in PVN feeding circuits..

**Figure 5.**
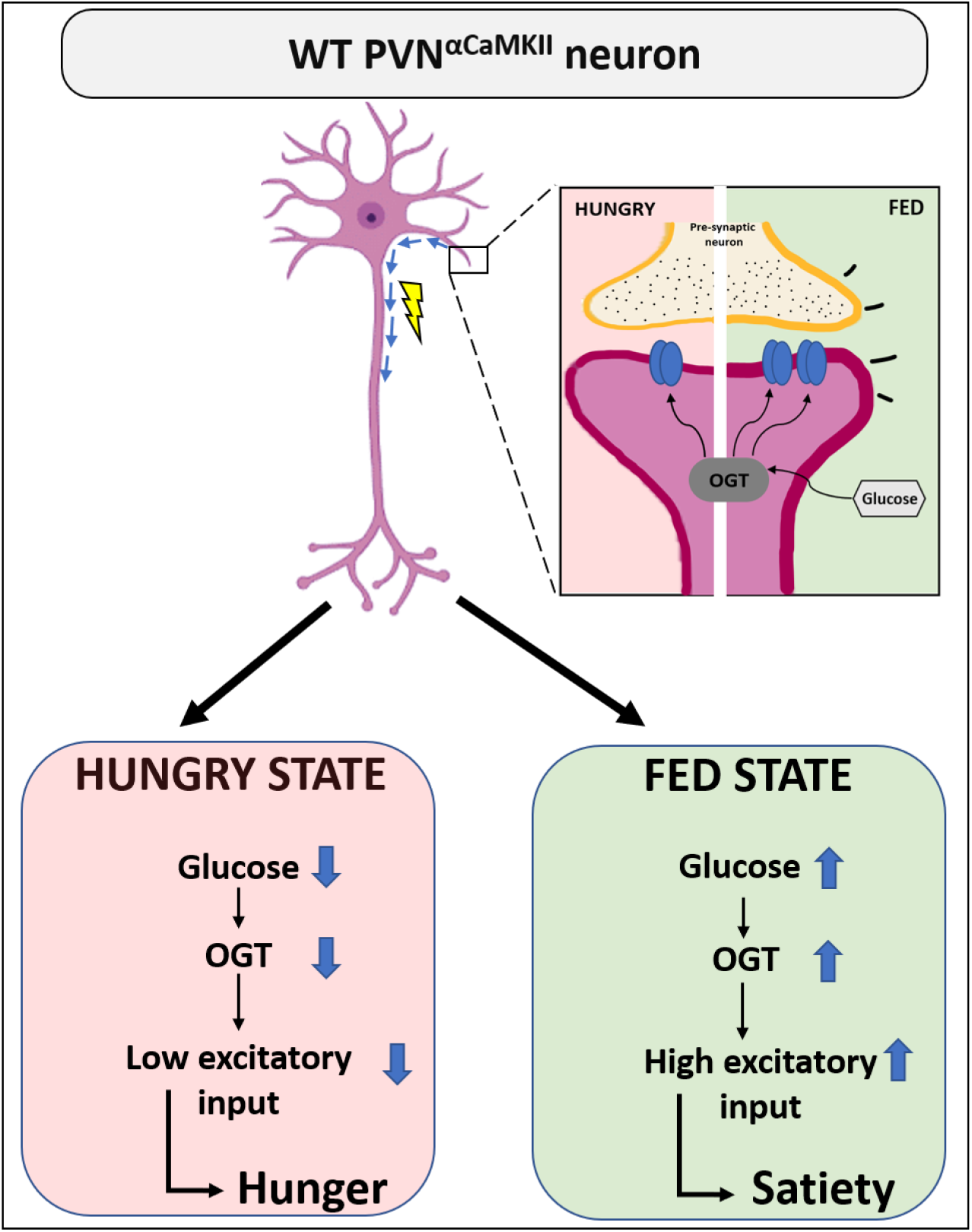
OGT functions as an adapter of excitatory synaptic plasticity in PVN^αCaMKII^ neurons across nutritional and feeding states. Schematic summarizing the role of OGT’s role as a postsynaptic adapter linking metabolic state to synaptic plasticity in feeding circuits across hungry (pink background) and fed (green background) states. **Top:** The enlarged inset depicts the synaptic interface where the pre-synaptic terminal (yellow) releasing neurotransmitters onto post-synaptic receptors in the PVN^αCaMKII^ neuron (magenta). OGT serves as a critical glucose sensor within the post-synaptic compartment, where glucose availability modulates OGT activity and subsequent O-GlcNAcylation of synaptic proteins, thereby controlling synaptic strength and stability. **Bottom panels**: In the hungry state (left), decreased glucose levels lead to reduced OGT activity, resulting in diminished excitatory synaptic input to promote hunger signaling. Conversely, in the fed state (right), elevated glucose enhances OGT function, increasing excitatory synaptic transmission and promoting satiation signaling.

## Discussion

This study demonstrates that OGT in PVN^αCaMKII^ satiation neurons adapts excitatory synaptic transmission to fluctuations in energy substrate availability. Deletion of OGT increases neuronal input resistance and excitability while preserving basic electrophysiological properties (Figure 1 and 2), and these intrinsic changes are accompanied by a disrupted ability to modulate excitatory synaptic input according to feeding status (Figure 3). Furthermore, OGT is necessary for synaptic responses to acute fluctuations in extracellular glucose, as evidenced by a pronounced synaptic depression in the OGT-KO neurons exposed to high glucose (Figure 4). These data suggest that OGT mediates satiation by coordinating neuron excitability and excitatory synaptic plasticity in response to nutrient availability. In contrast to the classical GLUT2/glucokinase/K_ATP_ system, our observations favor a model where OGT links satiation not only to intrinsic neuron excitability but makes satiation dependent on neuronal input from other brain regions to the PVN (Figure 1F and 1G). This model is further supported by our discovery that PVN^αCaMKII^ display several characteristics of neurons regulating metabolism by communicating with other neurons rather than with the neuroendocrine (Figure 1F and 1G). Importantly, by using low-titer viral stocks to achieve sparse OGT deletion, we were able to assess neuronal properties without confounding systemic effects; the lack of significant differences in body weight or daily average food intake between WT and OGT-KO animals (Figure 1D and E) confirms that our observations reflect cell-autonomous changes.

The basic membrane properties of PVN^αCaMKII^ neurons—resting membrane potential, AP threshold and capacitance—were not significantly altered by OGT deletion (Figure. 2C–E), indicating that OGT is not required for maintaining these baseline electrophysiological characteristics. However, deletion of OGT increased input resistance and lowered the excitability index (Figure.2F–G). Similarly, OGT has been shown to affect neuronal excitability in both AgRP neurons and the hippocampus[33, 34]. This indicates that OGT regulates how responsive individual neurons are to synaptic inputs[35].

The frequency and amplitude of excitatory inputs on PVN^αCaMKII^ turn out to be regulated by OGT and metabolic fluctuations. Spontaneous excitatory postsynaptic current (sEPSC) frequency increased nearly fourfold in wild-type neurons during feeding (Figure.3B), whereas this enhancement was absent in OGT-KO neurons (Figure.3F). Instead, in OGT-KO neurons, this feeding-dependent modulation is not only absent but there is a trend toward a paradoxical increased excitatory input in the hunger state is noted, although not statistically significant (Figure 3F). These data suggest that OGT is necessary for PVN^αCaMKII^ to adapt their excitatory synaptic input to changes in nutritional state[17, 24]. While increased excitability in OGT-KO neurons suggests they may intrinsically be more prone to firing, the effect on excitatory synaptic input indicate that OGT is necessary to couple their neuronal activity to metabolic fluctuations and thus mediate satiation upon eating. Indeed, the PVN^αCaMKII^ become activated upon food intake but this feeding-induced activation is blunted in OGT-KO neurons [17]. Nevertheless, our data indicate that is not the case that the PVN^αCaMKII^ neurons can no longer detect nutrients without OGT, rather OGT mediates satiation by enabling the neuron to adapt appropriately to nutritional fluctuations.

In addition to regulation of longer-term fluctuations of feeding state, our data show that OGT is necessary to preserve excitatory synaptic input during acute elevations in glucose (Figure 4.) Exposure to acute increases in extracellular glucose does not alter excitatory synaptic input in wild-type neurons(Figure 4B-D). However, in OGT-KO neurons, high glucose leads to significant reductions in both sEPSC frequency and amplitude (Figure 4F-G), representing maladaptive synaptic responses that inhibit satiation. Notably, decay tau is not significantly affected under any condition, highlighting that the observed effects are specific to the frequency and amplitude of synaptic events rather than kinetics (Figure 4D, H). Furthermore, there are corresponding differences in amplitude response fold-change (Figure 4J). This suggests OGT is critical for how excitatory synapses adapt to acute changes in nutrient availability.

The mechanism by which OGT adapts excitatory synaptic plasticity to metabolic fluctuations in PVN^αCaMKII^ may depend on AMPA receptor trafficking. It has been shown in the hippocampus that a rapid increase in O-GlcNAc levels through stimulating HBP flux decreases excitatory neurotransmission. This decrease is blunted if the major AMPA receptor subunit GluA2 is deleted [36]. We have shown also that AMPA receptor abundance in synapses is regulated by OGT [24]. The depression seen in OGT-KO neurons under high glucose (Figure 4F–G) could thus reflect disrupted AMPA receptor trafficking. Furthermore, O-GlcNAcase (OGA) activity likewise is shown to modulate AMPA receptor trafficking and dendritic spine morphogenesis, underscoring the dynamic interplay of OGT and OGA in regulating synaptic plasticity[37]. Much evidence indicates that AMPA receptor levels and neuronal activity in excitatory synapses can over time affect the number of functional synapses[38-41]. We have shown OGT regulates both excitatory synapse and dendritic spine number [24]. Thus, it is possible that OGT’s minutes-long effect on sEPSC frequency and amplitude upon acute elevation of glucose (Figure 4) can lead to the maladaptive hours-long change in sEPSC frequency dependent on feeding status (Figure 3). As we did not measure AMPA receptors directly in the PVN^αCaMKII^, transsynaptic factors may also contribute to the observed phenotype. We have shown previously that O-GlcNAc levels in PVN^αCaMKII^ are more sensitive to physiological, postprandial changes in glucose availability compared to other neurons[17]. Whereas there is a decrease in excitatory neurotransmission when deleting OGT in the PVN^αCaMKII^ as well as when stimulating OGT through the HBP in the hippocampus, these observations highlight a potential brain-region–specific difference in how OGT regulates excitatory neurotransmission depending on nutrient availability. Together, these results provide a coherent view where OGT enables PVN^αCaMKII^ neurons to mediate satiation by adapting excitatory synaptic input to both slow (feeding) and fast (glucose fluctuation) changes.

This mechanism supports a model in which OGT represents a fundamentally different class of nutrient sensor compared to canonical glucose-sensing mechanisms in the hypothalamus. The canonical GLUT2/glucokinase/K_ATP_ channel pathway primarily respond to glucose with an on-off effect on neuron excitability. While OGT also affect excitability, OGT’s regulation of synaptic plasticity in PVN^αCaMKII^ makes satiation directly dependent on input from upstream neurons. Anatomical mapping studies have reported that the PVN receives input from not only other hypothalamic nuclei but also, e.g., the amygdala, the bed nucleus of the stria terminalis (BNST) and the hippocampus [42-44]. Indirect input also comes from neocortex [45, 46]. This makes it possible that OGT regulates satiation by coupling metabolic fluctuations to emotional or cognitive aspects of food intake.

From a behavioral perspective, the feeding state-dependent regulation of excitatory synaptic transmission by OGT uncovers a mechanism by which OGT deletion in PVN^αCaMKII^ in adult mice leads to hyperphagia and impaired meal termination [17]. Importantly, OGT’s regulation of synaptic plasticity enables PVN^αCaMKII^ neurons to flexibly reconfigure and prioritize their upstream inputs according to the metabolic context—a functional capacity missing from classical glucose sensors. When OGT is knocked out, neuronal excitability is heightened; however, the inability to selectively filter and integrate incoming signals results in abnormal information processing and a breakdown in proper satiation signaling. This failure to appropriately up-regulate and stabilize excitatory signaling in response to nutritional cues directly inhibits satiation and leads to impaired meal termination [47]. Consequently, the disruption of energy intake regulation can manifest as either excessive consumption, contributing to obesity [5, 6, 48]. As stimulation of PVN^αCaMKII^ decreases food intake, it may in reverse also be associated with insufficient consumption, characteristic of anorexia nervosa[49], depending on the broader neural circuit context and the presence of compensatory mechanisms[50]. Taken together, these mechanisms provide a plausible foundation for understanding how metabolic dysfunction can contribute to unhealthy eating behaviors.

Overall, our data support a model in which OGT integrates metabolic information with synaptic machinery to ensure reliable excitatory neurotransmission in PVN^αCaMKII^ neurons across feeding and glucose fluctuations. This adaptive role provides a mechanistic basis towards understanding how the hypothalamus regulation satiation. These observations not only advance our mechanistic understanding of satiation signaling but also highlight how the brain integrates metabolic information to guide complex behavior, offering new perspectives on the neural basis of energy homeostasis and its dysregulation in disease.

## Materials and methods

### Animals and ethical approval of the study

Five adult male and five female C57BL/6J and OGT-KO mice were housed at the Umeå Center for Comparative Biology at 21 ± 2 °C, under a reversed light cycle (lights off at 23:00 h and on at 11:00 h). Food and water were provided ad libitum. All experimental procedures were approved and conducted in accordance with the regulations of the Local Animal Ethics Committee at Umea University.

Prior to conducting ex-vivo experiments, animals were equally divided into two groups: 10 mice in the 24-hour starved group and 10 in the non-starved group. The 24-hour starved animals were starved for 24 hours prior to perfusion, with free access to water. The non-starved animals had ad libitum access to food and water.

### Stereotactic virus injection

Anesthesia was induced using isoflurane gas inhalation, and anaesthetic depth was maintained at approximately 1.5% isoflurane. Anesthesia depth was monitored by regularly testing pinch reflexes and core temperature of the animal was maintained using a rectal probe. The coordinates for the PVN were as follows: distance from Bregma, −0.80 mm; distance from midline, ±0.20 mm; depth from surface, −4.80 mm. Animals were placed in a mouse stereotaxic head holder (WPI, UK). The skin was disinfected with Betadine, and a local anesthetic (Marcain® 2.5 mg/mL) was applied before incision to expose the skull. Bregma and Lambda were identified, and a cranial window was drilled over the injection site. The viruses pAAV-hSyn-DIO-EGFP (Addgene plasmid #50457; RRID:Addgene_50457) and pENN.AAV.CaMKII0.4.Cre.SV40 (Addgene plasmid #105558; RRID:Addgene_105558) were thawed, and a cocktail was created and diluted with sterile saline to two different final concentrations: 10x and 20x, respectively. The virus was then loaded into a pulled glass pipette (20-40 µm) and connected to a syringe pump (Nanoject III, Drummond Scientific). In total, bilateral injections of 500 nL each of the cocktail virus with 10x dilution were injected into 10 mice (5 WT and 5 OGT-KO), and 500 nL each of the 20x dilution were injected into another 10 mice (5 WT and 5 OGT-KO). The glass capillary was left in place for 5 minutes before being withdrawn. At the end of surgery, the skin was sutured, and the animals were given a subcutaneous injection of Carprofen (50 mg/mL) to relieve pain.

### Body weight and food intake assessment

Body weight and food intake were measured every alternate day after 10 days following stereotactic surgery until up to or maximum 6 days before tissue harvest. Mice were individually housed with free access to food and water. Body weight was recorded each morning using a precision balance. Food consumption was determined by weighing the food hopper at the same time each day and subtracting residual food weight to calculate daily intake. Data were averaged over the monitoring period for each animal with a maximum gap of 6 days before harvest. Statistical comparisons between WT and OGT-KO groups were performed using the Mann–Whitney non-parametric test, with significance set at P < 0.05.

### Immunohistochemistry

Mice were euthanized by cervical dislocation and immediately perfused intracardially with chilled artificial cerebrospinal fluid (aCSF). Serial brain sections were then cut using a microtome. Free-floating sections were blocked in blocking buffer (5% normal goat serum and 0.25% Triton X-100 in PBS) overnight at 4 °C on a shaker. Afterwards, they were incubated with primary antibodies overnight at 4 °C on a shaker, and subsequently with secondary antibodies for 2 hours on a shaker at room temperature. DAPI was included with the secondary antibodies. Primary and secondary antibodies were diluted in the blocking buffer. After both the primary and secondary antibody incubations, washes were performed using 0.25% Triton X-100 in PBS. The primary antibodies used were GFP (Thermo Fisher Scientific, cat. no. AB_2942817, dilution 1:1000) and OGT (Proteintech, cat. no. 11576-2-AP, dilution 1:1000). The secondary antibodies used were anti-GFP (Goat anti-chicken secondary antibody, AzureSpectra 490, cat. no. AC2209, dilution 1:1000) and anti-OGT (Goat anti-rabbit secondary antibody AzureSpectra 650, cat. no. AC2165, dilution 1:1000). After the final washes, sections were mounted on glass microscope slides and cover slipped using Invitrogen Fluoromount-G Mounting Medium. Slides were dried overnight at room temperature in the dark.

### Imaging analysis

Imaging was performed using a Leica SP8 confocal laser scanning microscope, and the same acquisition settings were maintained to analyse the paraventricular nucleus (PVN) in each WT and OGT-KO section. In the PVN, αCaMKII neurons were identified as cells expressing DAPI, GFP, and OGT signals in WT sections, and DAPI and GFP signals in OGT-KO sections. Images were processed and analysed using Fiji (ImageJ). The cell body diameter of each αCaMKII neuron was measured to classify them as either parvocellular or magnocellular neurons.

### Tissue preparation

Mice were euthanized by cervical dislocation without anaesthesia, followed by a quick transcardial perfusion with a Choline-Chloride (ChCl) based, ice-cold cutting solution, saturated with 100% oxygen containing (in mM): 110 ChCl, 26 NaAc, 2.5 Glucose, 11.6 Na-Ascorbate, 3.1 Na-Pyruvate, 10 HEPES, 2.5 KCl, 7 MgCl_2_, 0.5 CaCl_2_. This method was used to quickly cool down brain tissue and improve tissue survivability.

The brain was extracted and placed in the same ice-cold ChCl solution. Frontal and caudal ends of the brain were removed, and the remaining brain was placed in a 752M Vibroslicer (Campden Instruments Limited). Coronal slices of 300μm thickness containing PVN were obtained and incubated at 28°C in low glucose artificial cerebrospinal fluid (ACSF) saturated with 100% oxygen, containing (in mM): 125 NaCl, 2.5 KCl, 26 NaAc, 2 MgCl_2_, 2 CaCl_2_, 10 HEPES, 2.5 glucose, 17.5 sucrose.

### Electrophysiological recordings

PVN^αCaMKII^ neurons were identified via fluorescence using a monochromator controlled by TILLVisIon software (T.I.L.L. Photonics GmbH Munich, Germany). GFP-positive cells were selected for electrophysiological recordings. Conventional patch-clamp recordings were performed using borosilicate micropipettes (Harvard Apparatus, UK), made using a Flaming/Brown micropipette puller (Model P-97, Sutter Instrument CO, USA) to a resistance of 3-4 MΩ when submerged in the bath. Micropipettes were back-filled with a 0.2% biocytin intracellular solution containing (in mM): 110 K-gluconic acid, 10 NaCl, 1 MgCl_2_, 10 EGTA, 40 HEPES, 2 Mg-ATP, 0.3 Na-ATP.

Excitatory post-synaptic currents were recorded in voltage-clamp (VC) mode at a holding voltage of −53 to −65 m, and it was corrected for liquid junction potential. Liquid junction potential was calculated according to the stationary Nernst–Planck equation[51] using LJPcalc ([RRID:SCR_025044](https://scicrunch.org/resolver/SCR_025044)). Basic electrical properties of the neurons were measured in current-clamp (CC) mode, injecting currents in a sequential manner to detect the different thresholds for action potential and LTS, as well as to calculate neuron excitability.

Protocol began with CC recordings for Vm, AP threshold and LTS measurement, followed by a VC recording of sEPSCs for 5 min. Each set of protocols lasted approximately 10 min. The first two sets of recordings were performed under low (2.5 mM) glucose conditions. During the third VC recording, bath application of high (12.5 mM) glucose was initiated after one minute. The protocol continued in the same manner in high glucose for up to 30 minutes. Cells that survived the complete protocol were photographed and used to measure their size and compare with their respective capacitance to corroborate the selection of parvocellular neurons.

### Data analysis

Frequency, amplitude and decay tau were calculated from VC recordings in a semi-automated manner using MiniAnalysis software (Synaptosoft), whereas Vm, action potential threshold, resistance and LTS were calculated from CC files using Clampfit 11.2 (Molecular Devices, LLC.).

Statistics were performed using Origin 2021b (OriginLab Corporation) and GraphPad Prism 8. Mann-Whitney non-parametric test was used for comparing non-paired samples, while Wilcoxon signed rank test was used for paired samples. Statistical significance was set at a p-value<0.05.

Researchers involved in electrophysiological recordings and analysis were blinded during the data acquisition and analysis, revealing the genotype and feeding status of the animal only when the analysis was completed. This method was used to prevent any bias towards either condition, maintaining complete neutrality.

### Schematic representation

Figure 1A and figure 5 were generated using BioRender.com.

## References

1. Ahima RS, Antwi DA: Brain regulation of appetite and satiety. Endocrinology and metabolism clinics of North America 2008, 37(4):811–823.

2. Benelam B: Satiation, satiety and their effects on eating behaviour. Nutrition bulletin 2009, 34(2):126–173.

3. Woods SC: Gastrointestinal satiety signals I. An overview of gastrointestinal signals that influence food intake. American Journal of Physiology-Gastrointestinal and Liver Physiology 2004, 286(1):G7–G13.

4. Roger C, Lasbleiz A, Guye M, Dutour A, Gaborit B, Ranjeva J-P: The role of the human hypothalamus in food intake networks: An MRI perspective. Frontiers in nutrition 2022, 8:760914.

5. Timper K, Brüning JC: Hypothalamic circuits regulating appetite and energy homeostasis: pathways to obesity. Disease models & mechanisms 2017, 10(6):679–689.

6. Yeung A, Tadi P: Physiology, obesity neurohormonal appetite and satiety control.[Updated 2023 Jan 3]. StatPearls [Internet] Treasure Island (FL): StatPearls Publishing 2023.

7. Lin Z, Xuan Y, Zhang Y, Zhou Q, Qiu W: Hypothalamus and brainstem circuits in the regulation of glucose homeostasis. American Journal of Physiology-Endocrinology and Metabolism 2025, 328(4):E588–E598.

8. Fioramonti X, Chrétien C, Leloup C, Pénicaud L: Recent advances in the cellular and molecular mechanisms of hypothalamic neuronal glucose detection. Frontiers in physiology 2017, 8:875.

9. Miki T, Liss B, Minami K, Shiuchi T, Saraya A, Kashima Y, Horiuchi M, Ashcroft F, Minokoshi Y, Roeper J: ATP-sensitive K+ channels in the hypothalamus are essential for the maintenance of glucose homeostasis. Nature neuroscience 2001, 4(5):507–512.

10. Hussain S, Richardson E, Ma Y, Holton C, De Backer I, Buckley N, Dhillo W, Bewick G, Zhang S, Carling D: Glucokinase activity in the arcuate nucleus regulates glucose intake. The Journal of clinical investigation 2015, 125(1):337–349.

11. Evans ML, McCrimmon RJ, Flanagan DE, Keshavarz T, Fan X, McNay EC, Jacob RJ, Sherwin RS: Hypothalamic ATP-sensitive K+ channels play a key role in sensing hypoglycemia and triggering counterregulatory epinephrine and glucagon responses. Diabetes 2004, 53(10):2542–2551.

12. Lagerlöf O, Hart GW: O-GlcNAcylation of neuronal proteins: roles in neuronal functions and in neurodegeneration. Glycobiology of the Nervous System 2014:343–366.

13. Bond MR, Hanover JA: O-GlcNAc cycling: a link between metabolism and chronic disease. Annu Rev Nutr 2013, 33:205–229.

14. Speliotes EK, Willer CJ, Berndt SI, Monda KL, Thorleifsson G, Jackson AU, Allen HL, Lindgren CM, Luan Ja, Mägi R: Association analyses of 249,796 individuals reveal 18 new loci associated with body mass index. Nature genetics 2010, 42(11):937–948.

15. Gutierrez-Aguilar R, Kim DH, Woods SC, Seeley RJ: Expression of New loci associated with obesity in diet-induced obese rats: from genetics to physiology. Obesity 2012, 20(2):306–312.

16. Gonzalez-Rellan MJ, Fondevila MF, Dieguez C, Nogueiras R: O-GlcNAcylation: a sweet hub in the regulation of glucose metabolism in health and disease. Frontiers in endocrinology 2022, 13:873513.

17. Lagerlöf O, Slocomb JE, Hong I, Aponte Y, Blackshaw S, Hart GW, Huganir RL: The nutrient sensor OGT in PVN neurons regulates feeding. Science 2016, 351(6279):1293–1296.

18. Andersson B, Tan EP, McGreal SR, Apte U, Hanover JA, Slawson C, Lagerlöf O: O-GlcNAc cycling mediates energy balance by regulating caloric memory. Appetite 2021, 165:105320.

19. Krashes MJ, Shah BP, Madara JC, Olson DP, Strochlic DE, Garfield AS, Vong L, Pei H, Watabe-Uchida M, Uchida N: An excitatory paraventricular nucleus to AgRP neuron circuit that drives hunger. Nature 2014, 507(7491):238–242.

20. Zeltser LM, Seeley RJ, Tschöp MH: Synaptic plasticity in neuronal circuits regulating energy balance. Nat Neurosci 2012, 15(10):1336–1342.

21. Tolson KP, Gemelli T, Gautron L, Elmquist JK, Zinn AR, Kublaoui BM: Postnatal Sim1 deficiency causes hyperphagic obesity and reduced Mc4r and oxytocin expression. Journal of Neuroscience 2010, 30(10):3803–3812.

22. Lagerlöf O, Hart GW, Huganir RL: O-GlcNAc transferase regulates excitatory synapse maturity. Proceedings of the National Academy of Sciences of the United States of America 2017, 114(7):1684–1689.

23. Liu J, Conde K, Zhang P, Lilascharoen V, Xu Z, Lim BK, Seeley RJ, Zhu JJ, Scott MM, Pang ZP: Enhanced AMPA Receptor Trafficking Mediates the Anorexigenic Effect of Endogenous Glucagon-like Peptide-1 in the Paraventricular Hypothalamus. Neuron 2017, 96(4):897–909.e895.

24. Lagerlöf O, Hart GW, Huganir RL: O-GlcNAc transferase regulates excitatory synapse maturity. Proceedings of the National Academy of Sciences 2017, 114(7):1684–1689.

25. Fekete C, Lechan RM: Central regulation of hypothalamic-pituitary-thyroid axis under physiological and pathophysiological conditions. Endocr Rev 2014, 35(2):159–194.

26. Eliava M, Melchior M, Knobloch-Bollmann HS, Wahis J, da Silva Gouveia M, Tang Y, Ciobanu AC, Triana Del Rio R, Roth LC, Althammer F et al: A New Population of Parvocellular Oxytocin Neurons Controlling Magnocellular Neuron Activity and Inflammatory Pain Processing. Neuron 2016, 89(6):1291–1304.

27. Knobloch HS, Charlet A, Hoffmann LC, Eliava M, Khrulev S, Cetin AH, Osten P, Schwarz MK, Seeburg PH, Stoop R et al: Evoked axonal oxytocin release in the central amygdala attenuates fear response. Neuron 2012, 73(3):553–566.

28. Luther JA, Daftary SS, Boudaba C, Gould GC, Halmos KC, Tasker JG: Neurosecretory and non-neurosecretory parvocellular neurones of the hypothalamic paraventricular nucleus express distinct electrophysiological properties. J Neuroendocrinol 2002, 14(12):929–932.

29. Bean BP: The action potential in mammalian central neurons. Nature Reviews Neuroscience 2007, 8(6):451–465.

30. Zhang W, Linden DJ: The other side of the engram: experience-driven changes in neuronal intrinsic excitability. Nature Reviews Neuroscience 2003, 4(11):885–900.

31. Lazarus BD, Love DC, Hanover JA: O-GlcNAc cycling: implications for neurodegenerative disorders. The international journal of biochemistry & cell biology 2009, 41(11):2134–2146.

32. Dontaine J, Bouali A, Daussin F, Bultot L, Vertommen D, Martin M, Rathagirishnan R, Cuillerier A, Horman S, Beauloye C et al: The intra-mitochondrial O-GlcNAcylation system rapidly modulates OXPHOS function and ROS release in the heart. Commun Biol 2022, 5(1):349.

33. Ruan H-B, Dietrich MO, Liu Z-W, Zimmer MR, Li M-D, Singh JP, Zhang K, Yin R, Wu J, Horvath TL: O-GlcNAc transferase enables AgRP neurons to suppress browning of white fat. Cell 2014, 159(2):306–317.

34. Hwang H, Rhim H: Acutely elevated O-GlcNAcylation suppresses hippocampal activity by modulating both intrinsic and synaptic excitability factors. Scientific Reports 2019, 9(1):7287.

35. Yang YR, Song S, Hwang H, Jung JH, Kim SJ, Yoon S, Hur JH, Park JI, Lee C, Nam D et al: Memory and synaptic plasticity are impaired by dysregulated hippocampal O-GlcNAcylation. Sci Rep 2017, 7:44921.

36. Taylor EW, Wang K, Nelson AR, Bredemann TM, Fraser KB, Clinton SM, Puckett R, Marchase RB, Chatham JC, McMahon LL: O-GlcNAcylation of AMPA receptor GluA2 is associated with a novel form of long-term depression at hippocampal synapses. J Neurosci 2014, 34(1):10–21.

37. Han L, Galizia S, Pan J, Lagerlöf O: O-GlcNAcase promotes dendritic spine morphogenesis while downregulating their GluA2-containing AMPA receptors. bioRxiv 2025:2025.2008.2015.670533.

38. Kessels HW, Malinow R: Synaptic AMPA receptor plasticity and behavior. Neuron 2009, 61(3):340–350.

39. Shepherd JD, Huganir RL: The cell biology of synaptic plasticity: AMPA receptor trafficking. Annu Rev Cell Dev Biol 2007, 23(1):613–643.

40. Hill TC, Zito K: LTP-induced long-term stabilization of individual nascent dendritic spines. Journal of Neuroscience 2013, 33(2):678–686.

41. Mateos JM, Lüthi A, Savic N, Stierli B, Streit P, Gähwiler BH, McKinney RA: Synaptic modifications at the CA3–CA1 synapse after chronic AMPA receptor blockade in rat hippocampal slices. The Journal of physiology 2007, 581(1):129–138.

42. Li YJ, D. Wj, Liu R, Zan GY, Ye BL, Li Q, Sheng ZH, Yuan YW, Song YJ, Liu JG: Paraventricular nucleus-central amygdala oxytocinergic projection modulates pain-related anxiety-like behaviors in mice. CNS Neuroscience & Therapeutics 2023, 29(11):3493–3506.

43. Cole AB, Montgomery K, Bale TL, Thompson SM: What the hippocampus tells the HPA axis: Hippocampal output attenuates acute stress responses via disynaptic inhibition of CRF+ PVN neurons. Neurobiology of stress 2022, 20:100473.

44. Wang Y, Kim J, Schmit MB, Cho TS, Fang C, Cai H: A bed nucleus of stria terminalis microcircuit regulating inflammation-associated modulation of feeding. Nature communications 2019, 10(1):2769.

45. Liu Y, Li A, Bair-Marshall C, Xu H, Jee HJ, Zhu E, Sun M, Zhang Q, Lefevre A, Chen ZS et al: Oxytocin promotes prefrontal population activity via the PVN-PFC pathway to regulate pain. Neuron 2023, 111(11):1795–1811.e1797.

46. Land BB, Narayanan NS, Liu R-J, Gianessi CA, Brayton CE, M Grimaldi D, Sarhan M, Guarnieri DJ, Deisseroth K, Aghajanian GK: Medial prefrontal D1 dopamine neurons control food intake. Nature neuroscience 2014, 17(2):248–253.

47. Sternson SM, Eiselt AK: Three Pillars for the Neural Control of Appetite. Annu Rev Physiol 2017, 79:401–423.

48. Wang Q, Zhang B, Stutz B, Liu Z-W, Horvath TL, Yang X: Ventromedial hypothalamic OGT drives adipose tissue lipolysis and curbs obesity. Science Advances 2022, 8(35):eabn8092.

49. Frank GKW, Shott ME, DeGuzman MC: Recent advances in understanding anorexia nervosa. F1000Res 2019, 8.

50. Ghamari-Langroudi M, Srisai D, Cone RD: Multinodal regulation of the arcuate/paraventricular nucleus circuit by leptin. Proc Natl Acad Sci U S A 2011, 108(1):355–360.

51. Marino M, Misuri L, Brogioli D: A new open source software for the calculation of the liquid junction potential between two solutions according to the stationary Nernst-Planck equation. arXiv preprint arXiv:14033640 2014.

